# LSD persistently disrupts affective pain processing

**DOI:** 10.64898/2026.05.06.723205

**Authors:** Jared Plotkin, Elaine Zhu, Mélanie Druart, Qiaosheng Zhang, Eric Hu, Deven Cathcart, Nellie Jun, Leo Kwok, Tanya Sippy, Jing Wang

**Author notes:** These authors contributed equally (Jared Plotkin, Elaine Zhu, Mélanie Druart).

## Abstract

Psychedelics produce long-lasting effects, but their circuit mechanisms remain unclear. Here we show that, in rats, a single dose of lysergic acid diethylamide (LSD) persistently reduces pain affect. This effect is recapitulated by local administration in the anterior cingulate cortex (ACC), but not primary somatosensory cortex. Neuropixels recordings reveal that LSD suppresses stimulus-evoked nociceptive responses in the ACC, reducing the encoding of aversive value. Despite increasing intrinsic excitability ex vivo, LSD reduces the maximum stimulus-evoked firing of ACC neurons in vivo, indicating a dissociation between excitability and sensory encoding. Together, these findings show that psychedelics disrupt the cortical transformation of nociceptive input into aversive representations.

## Main

While pain has sensory and affective components, the affective symptoms often contribute strongly to patient suffering and comorbid psychiatric symptoms. Psychedelics have recently gained attention as potential therapeutic agents in clinical trials, where single-dose interventions have been shown to produce rapid and sustained improvements in affective symptoms^1–6^, with emerging evidence for effects on pain^8^. These results raise the possibility of long-term pain relief without the addiction liability associated with opioids.

Recent work demonstrates that psychedelics, including ketamine and psilocybin, can reverse pain-related behaviors and normalize elevated activity in the anterior cingulate cortex (ACC)^7,8^, a key hub for aversive processing within the cortex, especially in the context of nociceptive inputs^9–15^. Consistent with this, psilocybin has also been shown to reduce low-frequency oscillatory power and neuronal phase-locking in the ACC of awake rodents, suggesting a broader desynchronization of cortical activity^16^. However, these studies have primarily focused on changes in baseline or ongoing activity, leaving unresolved how psychedelics alter temporally specific nociceptive encoding. It also remains unclear whether these effects generalize beyond psilocybin to pharmacologically distinct agents such as lysergic acid diethylamide (LSD), which exhibits higher affinity and prolonged binding at the 5-HT2A receptor^17^. Notably, recent work reports limited or inconsistent analgesic effects of psilocybin across pain models, highlighting variability in sensory pain outcomes^18^. Together, these findings suggest that psychedelics may not uniformly alter nociceptive processing, but instead act on specific components of pain processing. In particular, it remains unknown whether psychedelics selectively reshape the transformation of nociceptive inputs into affective cortical representations^10,19^.

Here, we test the hypothesis that LSD reduces pain aversion by altering stimulus-evoked nociceptive encoding in the ACC. In rats, using a conditioned place aversion paradigm combined with in vivo electrophysiology, we examine whether LSD selectively reduces stimulus-evoked responses in ACC neurons while sparing sensory processing, and compare local effects in ACC and primary somatosensory cortex in regulating pain affect. We first examined the effects of systemic (intraperitoneal) LSD administration on the affective dimension of pain. To test the affective or aversive component of pain, we used a well-established conditioned place aversion assay (CPA)^20–23^. During a preconditioning phase, rats were allowed free access to two treatment chambers. During conditioning, one of the chambers was paired with repeated noxious stimulations of the hind paw (a pin-prick, or PP), and the opposite chamber was not paired with any noxious stimulations (no stimulus, or NS). Finally, during the testing phase, rats were given free access to both chambers without peripheral stimulations (**Fig. 1a**). Rats received intraperitoneal (i.p) saline or LSD (85 µg/kg) and were tested in the CPA paradigm 15 minutes later and again 14 days later without further drug exposure (**Fig. 1b**). This dose of LSD was sufficient to elicit the head twitch response, a rapid, stereotyped side-to-side head movement in rodents induced by hallucinogenic psychedelics^24–28^ (**Supplement 1a**). Importantly, LSD administration did not affect rat locomotion, as measured by distance traveled during either preconditioning (**Supplement 1b-c**) or testing (**Fig. 1c-e**; saline 3.1 ± .74 m, n = 12; LSD 3.8 ± m, n = 12; Student’s t-test: p = 0.22). As expected, rats given saline spent significantly less time in the PP chamber during testing versus preconditioning, indicative of baseline affective pain response (**Fig. 1f**; Preconditioning PP_preconditioning_ 315 ± 22.9 s, PP_testing_ 140 ± 25.5, n = 12, Student’s paired t-test: p=.0018). In contrast, after LSD administration, this aversive response to noxious stimulus was completely abolished (**Fig. 1g**; PP_preconditioning_ 325 ± 30.2; PP_testing_ 306 ± 44.7, n = 12; Student’s paired t-test: p=.74). The aversive response to pain was further quantified by a CPA score, calculated by subtracting the time rats stayed in the chamber paired with PP treatment during the testing phase from the preconditioning phase^11,12^. The CPA score was significantly larger for the saline than the LSD group (**Fig. 1h**; saline 175 ± 42.6, n = 12, LSD 19.2 ± 56.0, n = 12; Student’s t-test: p = 0.04). Impressively, when we repeated the CPA assays 14 days after this single dose of LSD administration, we found rats that were previously injected with saline again spent significantly less time in the PP chamber during testing versus preconditioning (**Fig. 1i**; PP_preconditioning_ 337 ± 33.0 s, PP_testing_ 35.4 ± 18.1, n = 6, Student’s paired t-test: p = 0.02) and that this effect remained attenuated in the rats previously treated with LSD (**Fig. 1j**; PP_preconditioning_ 294 ± 30.0 s, PP_testing_ 176.5 ± 92.63, n = 5, Student’s paired t-test: p=.016). The CPA score continued to be significantly larger in saline-than LSD-treated groups after 14 days (**Fig. 1k**; saline = 301.4 ± 43.0, n = 6; LSD = 118.1 ± 67.6, n = 5; Student’s t-test: p = 0.04). These results suggest that LSD produces a robust and enduring reduction in pain-associated aversion.

**Figure 1.**
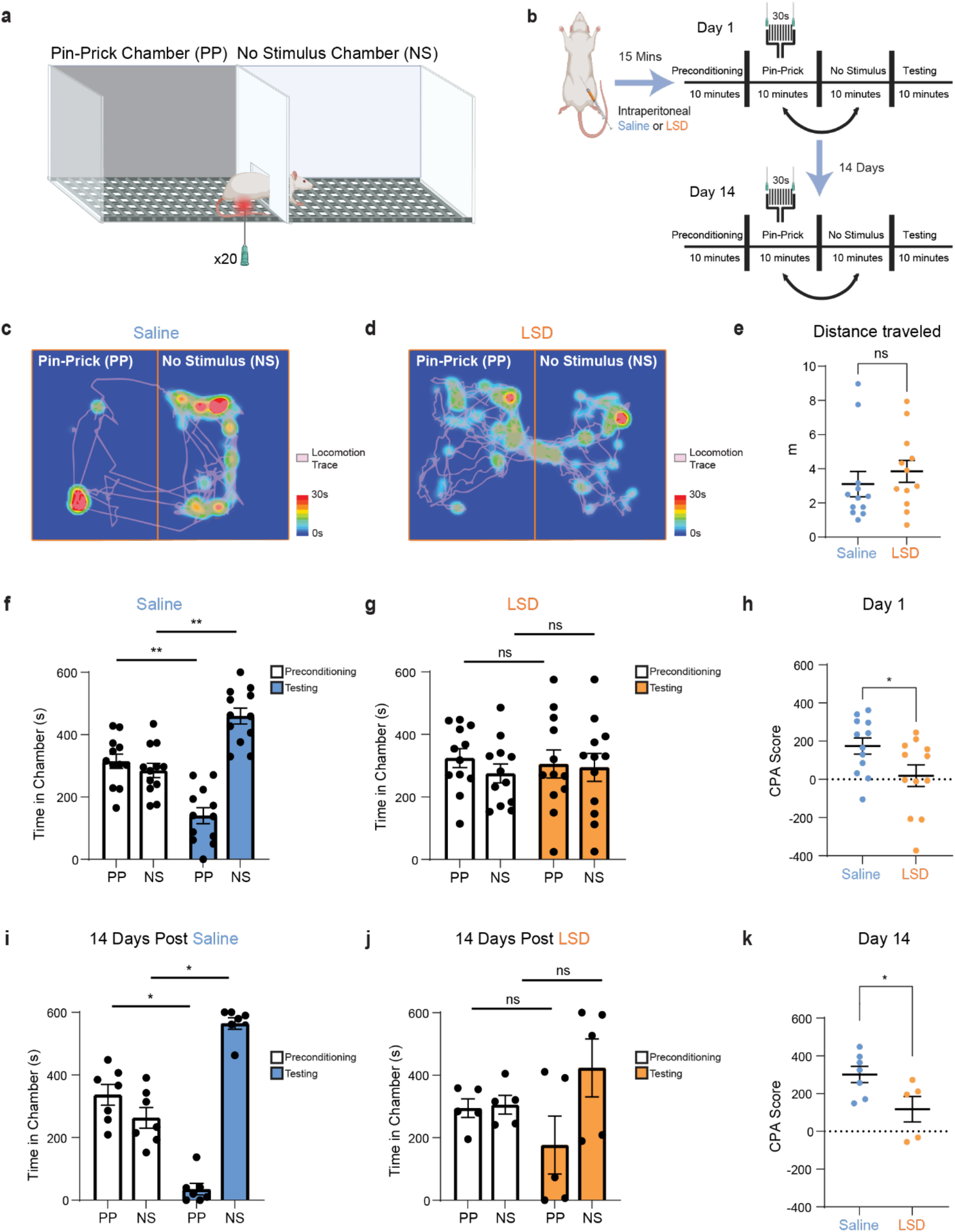
LSD produces immediate and persistent reductions in pain aversion. a. Schematic of conditioned place aversion (CPA) assay. Schematic created with BioRender.com. b. Timeline of behavioral experiments with IP injections. Schematic elements in this panel were created with BioRender.com. c. Locomotion heatmap of a rat given IP saline during testing phase. d. Locomotion heatmap of a rat given IP LSD during testing phase. e. LSD did not lead to any changes in rat locomotion during testing phase (unpaired Student’s t-test: ns p > 0.05, n=12) f. Rats given IP saline showed enhanced aversion to the pain-associated chamber (paired Student’s t-test: ** p < 0.01, n=12). g. Rats given IP LSD showed no aversion to pain-associated chamber (paired Student’s t-test: ns p > 0.05, n = 12). h. Rats given IP LSD (n = 12) showed less aversion to pain than rats given IP saline (n = 12) 15 min post-injection (unpaired Student’s t-test used: * p < 0.05). i. Rats given IP saline showed enhanced aversion to pain-associated chamber two weeks after injection (paired Student’s t-test: * p < 0.05, n=6). j. Rats given IP LSD showed no aversion to pain-associated chamber two weeks after injection (paired Student’s t-test: ns p > 0.05, n=5). k. Rats given IP LSD (n = 5) showed less aversion to pain than rats given IP saline (n = 6) two weeks post-injection (unpaired Student’s t-test: * p < 0.05).

The ACC is a well-known cortical region for regulating pain-aversive response^9–12,29^. To test if this brain area constitutes a target for the pain-relieving properties of LSD, we selectively infused 1 µL of LSD (1 mg/mL) into the ACC, followed by the CPA assay (**Fig. 2a-b**). We first confirmed that an equivalent volume of saline injected into the ACC preserved the aversive response to PP (**Fig. 2c**; PP_preconditioning_ 308 ± 30.0, PP_testing_ 131 ± 30.8, Student’s paired t-test: p=.0003). On the other hand, infusion of the LSD in the ACC blocked the aversive response to noxious inputs, as rats no longer avoided the chamber associated with PP (**Fig. 2d**; PP_preconditioning_ 310 ± 31.8, PP_testing_ 312 ± 70.7; Student’s paired t-test: p=.9736). The CPA score was significantly higher in saline than LSD infused animals (**Fig. 2e**; saline 177 +/- 30.9, n = 10, LSD -1.81 +/- 52.3, n = 6; Students t-test: p = 0.007). Similar to our results with IP injections, rats infused with saline showed aversion as quantified by time spent in PP chamber in preconditioning vs. testing periods (**Fig. 2f**; PP_preconditioning_ 358 ± 37.6, PP_testing_ 127 ± 60.1, n = 6; Student’s paired t-test: p = 0.03), while those infused with LSD did not show aversion (**Fig. 2g**; PP_preconditioning_ 260 ± 25.1, PP_testing_ 294 +/- 53.3, n = 6; Student’s paired t-test: p = 0.60). The CPA scores 14 days after infusion into the ACC were also significantly different between control and LSD (**Fig. 2h**; saline 231 ± 69.5, n = 6; LSD -34.6 ± 58.9, n = 6; Mann-Whitney U test: p = 0.02). The amplitude and time course of these effects are similar to those produced by systemic LSD administration, suggesting that the ACC is sufficient to mediate these pain-related effects.

**Figure 2.**
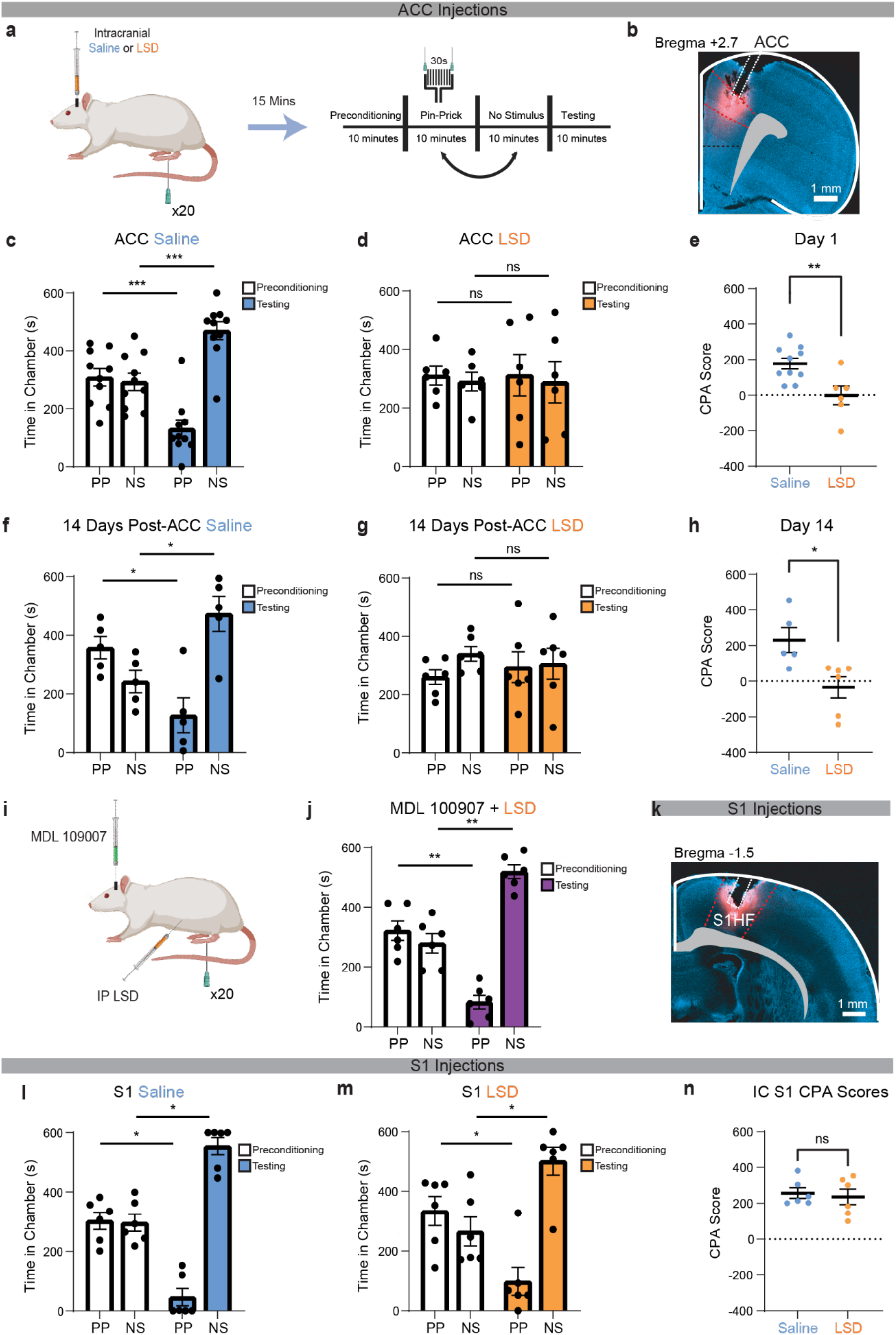
LSD disrupts pain aversion via 5-HT2A receptors specifically in the ACC. a. Timeline of behavioral experiments involving bilateral intracranial injections. Schematic elements in this panel were created with BioRender.com b. Histological image showing verification of ACC injection site by labeling with the fluorescent dye, BODIPY TMR-X. c. Rats given intracranial saline into the ACC showed enhanced aversion to the pain-associated chamber (paired Student’s t-test: *** p < 0.001 used, n=10). d. Rats given intracranial LSD into the ACC did not show aversion to pain-associated chamber (paired Student’s t-test: ns p > 0.05, n=6). e. Rats given intracranial LSD (n = 6) showed less aversion to pain than rats given intracranial saline (n = 10) 15 min post-injection (unpaired Student’s t-test: **p < 0.01). f. Rats given intracranial saline into the ACC showed enhanced aversion to pain-associated chamber two weeks after injection (paired Student’s t-test: * p < 0.05, n=6). g. Rats given intracranial LSD into the ACC showed no aversion to pain-associated chamber two weeks after injection (paired Student’s t-test: ns p > 0.05, n=6). h. Rats infused with LSD into the ACC (n = 6) showed less aversion to pain than rats infused with saline into the ACC (n = 6) two weeks post-injection (unpaired Student’s t-test used: * p < 0.05). i. Schematic of injection strategy. MDL 100907 was injected bilaterally into ACC 15 min before LSD was injected IP. Rats underwent the CPA assay 15 minutes after LSD injection. Schematic created with BioRender.com. j. Rats given intracranial MDL 100907 + IP LSD show enhanced aversion to the pain-associated chamber (paired Student’s t-test: ** p < 0.01, n=6) k. Histological image showing verification of S1 injection site by labeling with the fluorescent dye, BODIPY TMR-X. l. Rats that were given intracranial saline into S1 showed enhanced aversion to the pain-associated chamber (Wilcoxon signed rank test: * p < 0.05, n=6). Schematic created with BioRender.com m. Rats that were given intracranial LSD into S1 showed enhanced aversion to the pain-associated chamber (Wilcoxon signed rank test: * p < 0.05, n=6). n. Rats given intracranial LSD into S1 showed no difference in aversion to pain than rats given intracranial saline (unpaired Student’s t-test: ns p > 0.05).

To further elucidate the mechanism through which LSD exerts this effect, we explored how targeted pharmacological blockage of the 5-HT_2A_ receptor with MDL 100,907 (MDL) in the ACC would modulate the affective pain response in rats administered IP LSD in the CPA assay (**Fig. 2i**). We found that MDL significantly attenuated the anti-aversive effects of the LSD, as rats now avoided the PP chamber (**Fig. 2j**; PP_preconditioning_ 321 ± 32.3, PP_testing_ 82.1 ± 22.8, n = 6; Student’s paired t-test: p=.001) As a further comparison, and to evaluate the sensory aspects of pain, we infused saline and LSD directly into the primary hindlimb somatosensory cortex (S1), a cortical region that is important for the sensory but not the affective component of pain^30^ (**Fig. 2k**). Rats infused with saline in S1 showed significant aversion, as quantified by time spent in PP chamber in preconditioning versus testing (**Fig. 2l**; PP_preconditioning_ 303 ± 70.2, PP_testing_ 45.9 ± 71.3, n = 6; Wilcoxon signed rank test: p = 0.03), as did those injected with LSD in S1 (**Fig. 2m**; PP_preconditioning_ 334 +/- 119, PP_testing_ 98.5 +/- 116, n = 6; Wilcoxon signed rank test: p = 0.03). The CPA score was not different between saline and LSD groups (**Fig. 2n;** Saline 257 ± 72.4; LSD 236 ± 106; Student’s unpaired t-test: p=.6953). These results support the specificity of the ACC in mediating the pain-relieving effects of LSD.

To examine the effects of LSD on neuronal activity in the ACC, we used chronically implanted Neuropixels 2.0 probes to record spiking activity in freely behaving rats following intraperitoneal infusion of LSD or saline. In these experiments, rats received the injection and were then subjected to hindpaw pin-prick (PP) stimulation during recordings, independent of the CPA paradigm (**Fig. 3a**). Probe placement in the ACC was confirmed by post hoc histology (**Fig. 3b**). Consistent with previous reports^12,21^, a fraction of neurons increased their firing rates in response to PP, and this proportion, while fewer in LSD treated rats, was not significantly different (**Fig. 3c-d; Supplementary Fig. 3a-b**). Neurons recorded from rats that received LSD, however, exhibited lower peak firing rates in response to PP compared to those recorded from saline-treated rats (**Fig. 3e-g, Supplementary Fig. 2c;** Saline 6.44 ± 0.40 Hz, n = 515, LSD 5.01 ± 0.32, n = 486; Mann-Whitney U test: p = 0.006). This reduction persisted and remained significant 14 days after LSD administration (**Fig. 3h, Supplementary Fig. 2d, 3a;** saline 5.38 ± 0.32, n = 569, LSD 4.25 ± 0.34, n = 372; mean ± SEM, Mann-Whitney U test: p=.0131). We next focused on pain-responsive neurons, defined as neurons that significantly increased their firing rates in response to the noxious PP stimulus (see Methods). Among these neurons, LSD also reduced nociceptive responses compared to saline (**Fig. 3i–k, Supplementary Fig. 2c**; saline 15.0 ± 1.22 Hz, n = 120; LSD 10.5 ± 1.00 Hz, n = 90; Mann–Whitney U test, *p* = 0.006). This decrease in peak firing rate persisted 14 days after LSD administration and remained significant (**Fig. 3l, Supplementary Fig. 2f, 3b**; saline 14.4 ± 1.52 Hz, n = 85; LSD 10.5 ± 2.22 Hz, n = 42; Mann–Whitney U test, *p* = 0.005).

**Figure 3.**
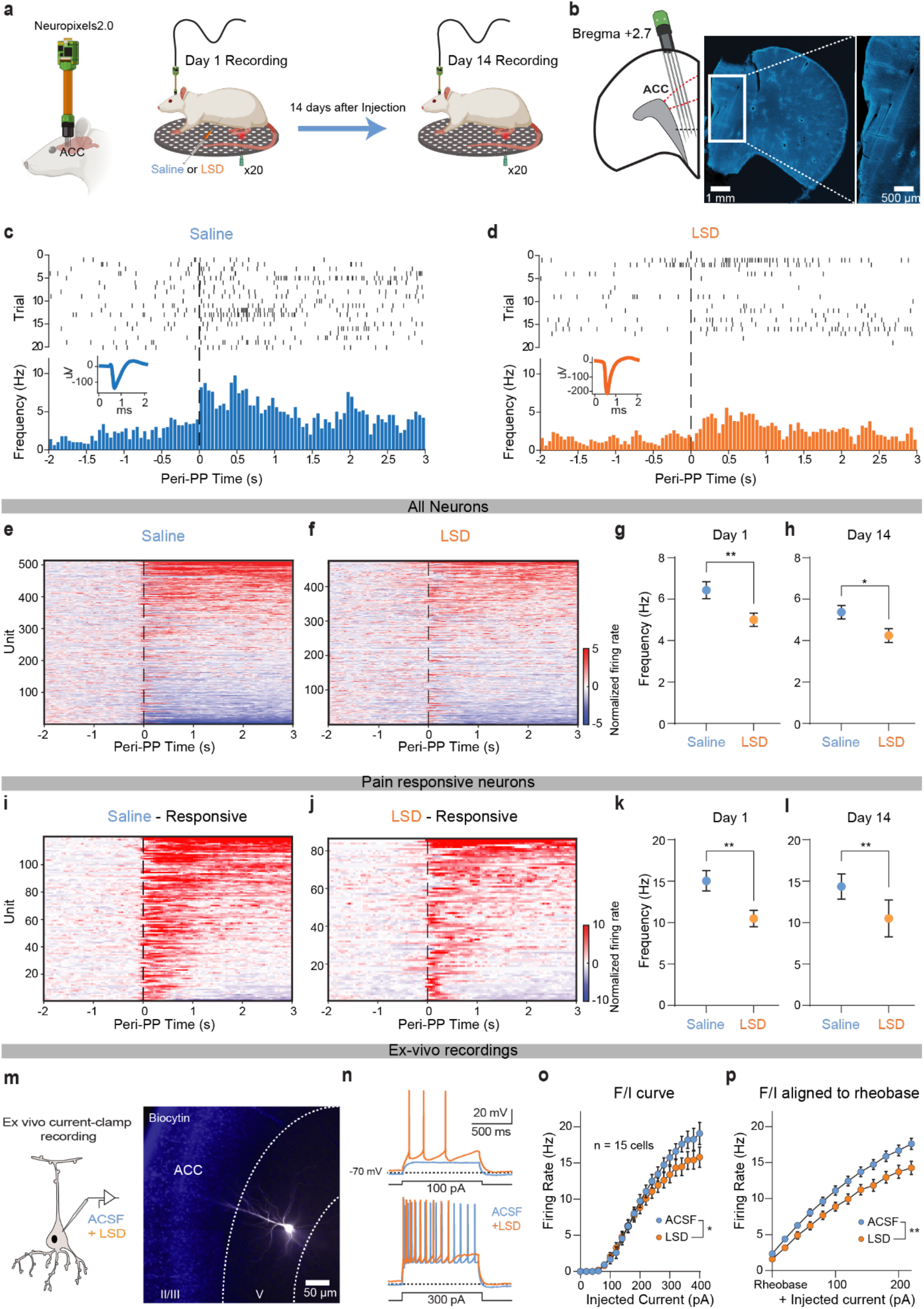
LSD persistently reduces stimulus-evoked nociceptive responses in the ACC. a. Schematic of experiment. Rats were injected with either LSD or saline and then probed with PP stimulus on day 1, 15 minutes after injection, and day 14. Schematic created with BioRender.com b. Left: schematic of the Neuropixels 2.0 probe recording site. Right: representative histological image confirming probe placement. Schematic elements in this panel were created with BioRender.com c. Raster plot and peristimulus time histogram (PSTH) of a neuron following saline exposure. d. Raster plot and peristimulus time histogram (PSTH) of a neuron following LSD exposure. e. Heatmap of neuronal activity following saline exposure on day 1. Pin-prick applied at time 0; firing rates are normalized to the pre–pin-prick baseline. n = 515 neurons from 4 rats. f. Heatmap of neuronal activity following LSD exposure on day 1. Pin-prick applied at time 0; firing rates are normalized to the pre–pin-prick baseline. n = 486 neurons from 4 rats. g. ACC neurons recorded from rats exposed to LSD show reduced baseline-subtracted peak firing rates in response to pin-prick on day 1 of injection. (unpaired Student’s t-test: ** p < 0.01). h. ACC neurons recorded from rats exposed to LSD show reduced baseline-subtracted peak firing rates in response to pin-prick on day 14 of injection. (unpaired Student’s t-test: * p < 0.01). i. Heatmap of pain-responsive neurons following saline exposure on day 1. Pin-prick applied at time 0; firing rates are normalized to the pre–pin-prick baseline. n = 120 neurons from 4 rats. j. Heatmap of pain-responsive neurons following LSD exposure on day 1. Pin-prick applied at time 0; firing rates are normalized to the pre–pin-prick baseline. n = 90 neurons from 4 rats. k. ACC pain-responsive neurons exposed to LSD exhibit reduced baseline-subtracted peak firing rates in response to pin-prick on day 1 (unpaired Student’s t-test, **p < 0.01). l. ACC pain-responsive neurons exposed to LSD exhibit reduced baseline-subtracted peak firing rates in response to pin-prick on day 14 (unpaired Student’s t-test, **p < 0.01). m. Left: schematic of an ACC layer 5 pyramidal neuron recorded in acute slice using whole-cell patch-clamp. Right: histological reconstruction of a biocytin-filled neuron. n. Example action potential (AP) trains evoked by somatic current injections of 100 pA (top) and 300 pA (bottom) before (blue) and after incubation with LSD (orange). o. F–I curve for ACC pyramidal neurons showing decreased excitability following LSD incubation (two-way ANOVA: *p < 0.05, n = 15 cells from 6 rats). p. F–I relationship normalized to the rheobase of each condition (mean ± s.e.m.), highlighting a reduction in intrinsic excitability during LSD perfusion (two-way ANOVA: **p < 0.01, n = 15 cells from 6 rats).

To examine how LSD affects intrinsic firing properties at the cellular level, we performed ex vivo current-clamp recordings from ACC pyramidal neurons in layer 5 (**Fig. 3m**). Neurons exposed to LSD fired more readily at low current injections (**Fig. 3n**; **Supplementary Fig. 3c**). However, at higher current injections, firing rates declined and were significantly lower than those of neurons recorded in ACSF alone (**Fig. 3o**; two-way repeated-measures ANOVA, drug effect: F(1,14) = 6.41, *p* = 0.024; n = 15 neurons from 6 rats). This effect was more pronounced when firing rates were aligned to rheobase (**Fig. 3p**; two-way repeated-measures ANOVA, drug effect: F(1,14) = 12.91, *p* = 0.003). These findings are consistent with our Neuropixels recordings, which show reduced peak firing in vivo in response to pain stimuli. Together, the results indicate that LSD increases intrinsic excitability while constraining maximal firing output, providing a potential reconciliation of prior findings that psychedelics enhance neuronal excitability in slice preparations yet reduce firing rates in vivo.

In our study, a single exposure to LSD produces a persistent reduction in the affective component of pain, accompanied by a selective but sustained suppression of stimulus-evoked nociceptive responses in the anterior cingulate cortex (ACC). These effects are regionally specific, as local ACC, but not primary somatosensory cortex, administration recapitulates the behavioral phenotype. While recent studies have shown that psilocybin can normalize elevated activity in ACC circuits, our results demonstrate that psychedelics selectively alter stimulus-evoked nociceptive encoding, rather than simply modulating overall levels of cortical activity. Moreover, as our findings extend these observations from ketamine and psilocybin to LSD, they highlight a robust modulating effect on the affective, rather than sensory, dimension of pain that may be a generalizable feature of psychedelics.

At the cellular level, LSD increases intrinsic excitability of ACC pyramidal neurons while constraining maximal firing output, providing a potential explanation for the reduced peak stimulus-evoked responses observed in vivo. This dissociation helps reconcile prior findings that psychedelics enhance neuronal excitability^31,32^ and excitatory neurotransmission^33,34^ in slice preparations yet reduce firing rates in intact circuits^35–37^. Together, these results support a model in which psychedelics disrupt the transformation of sensory input into aversive cortical representations. Future studies can expand from our findings to further define the contributions of specific neuronal subtypes and local microcircuit dynamics within the ACC.

## Methods

### Experimental protocol and data acquisition

All experimental studies were conducted in accordance with the New York University School of Medicine (NYUSOM) Institutional Animal Care and Use Committee (IACUC) regulations to ensure minimal animal use and discomfort, license reference number: IA16-01388. Male Sprague– Dawley rats were purchased from Taconic Farms and kept in a rearing room facility in the NYU Langone Science Building, controlled for humidity, temperature, and a 12-h (6:30 a.m. to 6:30 p.m.) light–dark cycle. Food and water were available ad libitum. Animals arrived at the facility weighing 250 to 300 g and had an average of 10 days to acclimate to the new environment before the experiment began. Experiments involving the use of LSD were conducted under appropriate DEA licensure

### CPA assay

Rats were injected either intraperitoneally or intracranially 15 minutes before they undergo experimentation. They spent this 15 minutes in their home cage. At the beginning of the assay, the rat was placed in a two-chamber apparatus consisting of equally sized compartments, which were connected by a large opening that allowed free movement between the chambers. A different scented balm was applied to the walls of each chamber to provide the rat with contextual cues. The behavioral paradigm consisted of preconditioning (baseline), conditioning, and testing phases. During the preconditioning phase (10 min), the rat was allowed to roam freely between the two chambers without any stimulus from the experimenter. Animals that spent more than 480 s or less than 120 s of the total time in either chamber (>80% preference) during this phase were eliminated from further analysis. Immediately after the preconditioning phase, the opening between the two chambers was closed. The rat would then undergo conditioning. For ten minutes the rat could roam around one half of the two chamber apparatus and no stimulus was applied. For the next ten minutes the rat was moved to the other half of the two chamber apparatus and a noxious pin-prick stimulation with a 27G needle was applied to the plantar surface of the right hind-paw every 30 seconds. The order of the pin-prick stimulus and the no stimulus were counterbalanced, such that half of the rats received the pin-prick stimulation first, while the other half received no stimulus first during conditioning. Chamber pairings were also counterbalanced. During the testing phase (10 min), the barrier between the two chambers was opened up again. The rat was placed in the center of the apparatus and no stimulation was given by the experimenter. The rat was allowed to travel freely between the two chambers. Rats which moved less than 0.5m during the testing phase were excluded. AnyMaze software and a video camera were used to track the movements of the rat in each chamber. Decreased time spent in a chamber during the testing phase compared to the preconditioning phase indicated avoidance (aversion) of that chamber, while increased time in a chamber indicated a preference for that chamber. The CPA score, which quantifies an animal’s aversion to the stimulus, was computed by subtracting the time the rat spent in the chamber associated with the pin-prick stimulation during the tesing phase from the time it spent in the same chamber during the preconditioning phase. A higher CPA score indicated greater aversion to the pin-prick stimulus.

### Cannula implantation and intracranial injections

Rats were anesthetized with 1.5–2% isoflurane and 8 mm 15G steel guide cannulas (Protech International Inc.) were bilaterally inserted. When targeting the ACC these cannulas were inserted at anteroposterior (AP) + 2.7 mm, mediolateral (ML) ± 1.6 mm, and dorsoventral (DV) -1 mm at an angle of 17° toward the midline. When targeting S1 these cannulas were inserted at anteroposterior (AP) - 1.5 mm, mediolateral (ML) ± 2.5 mm, and dorsoventral (DV) -0.5 mm at an angle of 10° toward the midline. Dental acrylic was used to keep the guide cannulas in place. After insertion rats were placed on a heating pad until they recovered from anesthesia and were monitored daily. Rats were allowed one week to recover post-cannula implantation before injections were done. For injections, rats were anesthetized with 1.5–2% isoflurane. For intracranial injections of LSD, saline, or MDL 100,907, 1 microliter of solutions were loaded into two 30 cm lengths of PE-50 tubing attached at one end to 10 μl Hamilton syringes filled with distilled water and at the other end to 33 gauge injector cannulas, which extended 1 mm beyond the implanted guides for the ACC. Injections were done over 100s and then the injector cannula remained in the guide for an additional 60s and slowly raised. Behavioral experiments occurred 15 minutes after rats awoke from anesthesia.

### LSD Dosage

IP injections are done at a dose of 85 micrograms/kg. LSD, sourced from National Institute on Drug Abuse Drug Supply Program (NDSP, 95% purity) was taken from aliquots dissolved in saline with a concentration of 0.5 mg/kg. Equivalent volume of saline is injected as a control. For intracranial injections one microliter is injected into each hemisphere. Intracranial aliquots are 1 mg/kg. MDL 100,907 intracranial injections aliquots are 1 microgram/mL and 1 microliter was injected into each hemisphere.

### Neuropixels implantation

We used Neuropixels 2.0 probes (NP2014) to record multichannel neural activities from the rat ACC, on the contralateral side of the paw that received noxious stimulation. Neuropixels probes were glued with 3D printed custom design drives before implantation. For surgery, rats were anesthetized with isoflurane (1.5%-2%). The skull was exposed, and a 2 mm-diameter hole was drilled above the target region. The coordinates for the ACC implants were AP 2.9, ML 1-1.7, DV 5.0, with an angle of 17° toward the midline. After insertion, the craniotomy was covered with silicone artificial dura gel (Cambridge NeuroTech) to protect the dura. Vaseline was used to wrap probe movable parts, including the probe shanks and flexible cables, as well as the drive shuttle. Ground and reference wires were inserted separately into burr holes in the cerebellum. Both drive and probe connected with 2.0 Headstage (HS_2010) were secured to the skull screws using dental cement. After surgery, rats were placed on a heating pad until they recovered from anesthesia and were monitored daily. Rats were allowed at least one week to recover post-implantation before IP injections and behavioral neural recordings were done.

### Neuropixels behavioral assay

The rat is injected IP with either LSD or saline and then placed in their home cage for 15 minutes. The rat is then placed in a chamber and neuropixels recording begins. Sessions are recorded with a 30 fps camera from below to align time of pin-prick to the neural recording. The rat is allowed to freely roam the chamber for 5 minutes. 20 Pin-pricks are then applied to the hindpaw contralateral to the implanted neuropixels every minute. Two weeks after the injection this paradigm is repeated without a second injection.

### Electrophysiological recordings and spike-sorting

Neural signals were recorded through a custom PXIe acquisition module via a PXI chassis (NI 1071) or Neuropixels OneBox system (ONEBOX_1000). Signals from ACC area are collected at 30 kHz using SpikeGLX software. This was then spike-sorted through Kilosort4. These results are manually adjusted using Phy2. To ensure no bias in manual curation, initial recordings were separately sorted by two lab members. The quantitative results yielded from each of these sorted files were similar and led to the same statistical conclusions.

### Data analysis

This study examines the peak firing rate of neurons following pin-prick. To determine peak firing rate, for each neuron the maximum firing rate within 5 s after each pin-prick is calculated. This is averaged across all 20 trials of pin-pricks to determine peak firing rate for each neuron. Peri-stimulus firing rates were computed using 50 ms bins, followed by smoothing with a 250 ms moving average filter. Next, we computed the Z-scored firing rates using each neuron’s own activity during baseline (defined as its distribution of firing rates within a 3 min habituation period at the start of the recording, using 1 s bins, smoothed with a 2.5 s moving average filter). A neuron was called a positive pain-responding neuron if the absolute value of the Z-scored firing rate of at least four consecutive time bins (total 200 ms) after stimulation was greater than 2.6 (two-tailed p < 0.005)^38^. This criteria must be fulfilled within 1 s after the stimulus onset.

### Histology

Histology was performed to verify location of intracranial injections and neuropixels probe placement. For intracranial injections, 1 microliter of fluorescent dye (BODIPY TMR-X) was injected into each hemisphere using the already placed cannulas. This is done to observe the spread and location of the injection. Rats were then deeply anesthetized using isoflurane (1.5%-2%) and transcardially perfused with phosphate buffered saline (1 minute) and then paraformaldehyde (9 minutes). The brain was removed and stored at 4° C in 4% paraformaldehyde for 48 hours. After 48 hours the paraformaldehyde is replaced with phosphate buffered saline. Coronal 100 micrometer slices were then collected and mounted with DAPI.

### Acute slice preparation and ex vivo electrophysiology

Acute brain slices were prepared from naïve rats. Animals were deeply anesthetized with isoflurane (5%) and transcardially perfused with ice-cold (0–4 °C), oxygenated cutting solution containing (in mM): 110 choline chloride, 2.5 KCl, 25 glucose, 25 NaHCO_3_, 1.25 NaH_2_PO_4_, 0.5 CaCl_2_, 7 MgCl_2_, 11.6 L-ascorbic acid, and 3.1 sodium pyruvate, continuously bubbled with 95% O_2_/5% CO_2_. Brains were rapidly extracted, and 300 µm-thick coronal slices were cut in the same solution using a vibratome (Leica VT1200S).

Slices were transferred to artificial cerebrospinal fluid (ACSF) containing (in mM): 125 NaCl, 2.5 KCl, 25 glucose, 25 NaHCO_3_, 1.25 NaH_2_PO_4_, 2 CaCl_2_, and 1 MgCl_2_, continuously bubbled with 95% O_2_/5% CO_2_. Slices were incubated at 32 °C for 20 min and subsequently maintained at room temperature (20–25 °C) until recording. For electrophysiological recordings, slices were transferred to a submerged recording chamber and continuously superfused with oxygenated ACSF at 30–32 °C.

Whole-cell current-clamp recordings were obtained from layer V pyramidal neurons in the ACC using borosilicate pipettes (3–5 MΩ) filled with potassium-based intracellular solution containing the following (in mM): 135 K, 5 KCl, 0.1 EGTA-KOH, 10 HEPES, 2 NaCl, 5 MgATP, 0.4 Na2GTP, 10 Na2-phosphocreatine and 2–4 mg/mL of biocytin. Intrinsic properties and neuronal excitability were assessed by generating spike frequency–current (F–I curve) using a series of 1s depolarizing current steps incremented by 25 pA. After baseline recordings, LSD (700 nM) was bath-applied, and F–I curves were reassessed after 5–10 min.

### Statistics

Data are presented as mean ± SEM. Statistical analyses were performed using Prism (Graphpad). The normality of data distribution was tested using Shapiro-Wilk’s test. Unpaired two-tailed t-tests (for normally distributed datasets) or Mann–Whitney tests (for non-normally distributed datasets) were used for comparisons between two groups. For CPA results within groups, paired two-tailed t-tests (for normally distributed datasets) or Wilcoxon tests (for non-normally distributed datasets) were used. Values of P < 0.05 were considered statistically significant.

## Acknowledgements

We thank the entire Sippy and Wang Labs for helpful discussions, Corryn Chaimowitz and Sarah Mennenga for technical assistance, and the NIDA Drug Supply Program for supplying LSD. This work was supported by NIH grants R01 NS126391 (T.S.), R01 5R01MH130658 (T.S.), R01 GM115384, T32 GM136573 (J.P.), Leon Levy Foundation Scholar Award (M.D) and the NYU Interdisciplinary Pain Research Program (J.W.).

## Data availability

The main data supporting the findings of this study are available within the paper and its Supplementary Information. At the time of publication, all datasets will be deposited in Zenodo.

## Code availability

The custom code used in this study is available on GitHub.

## References

1. Carhart-Harris, R. et al. Trial of Psilocybin versus Escitalopram for Depression. N Engl J Med 384, 1402–1411 (2021).

2. Carhart-Harris, R. L. et al. Psilocybin with psychological support for treatment-resistant depression: an open-label feasibility study. The lancet. Psychiatry 3, 619–27 (2016).

3. Davis, A. K. et al. Effects of Psilocybin-Assisted Therapy on Major Depressive Disorder: A Randomized Clinical Trial. JAMA Psychiatry 78, 481–489 (2021).

4. Griffiths, R. R. et al. Psilocybin produces substantial and sustained decreases in depression and anxiety in patients with life-threatening cancer: A randomized double-blind trial. J Psychopharmacol 30, 1181–1197 (2016).

5. Ross, S. et al. Rapid and sustained symptom reduction following psilocybin treatment for anxiety and depression in patients with life-threatening cancer: a randomized controlled trial. J Psychopharmacol 30, 1165–1180 (2016).

6. Goodwin, G. M. et al. Psilocybin for treatment resistant depression in patients taking a concomitant SSRI medication. Neuropsychopharmacol. 48, 1492–1499 (2023).

7. Hammo, A., Wisser, S. & Cichon, J. Single-dose psilocybin rapidly and sustainably relieves allodynia and anxiodepressive-like behaviors in mouse models of chronic pain. Nat Neurosci 28, 2285–2295 (2025).

8. Zhou, H. et al. Ketamine reduces aversion in rodent pain models by suppressing hyperactivity of the anterior cingulate cortex. Nat Commun 9, 3751 (2018).

9. Rainville, P., Duncan, G. H., Price, D. D., Carrier, B. & Bushnell, M. C. Pain affect encoded in human anterior cingulate but not somatosensory cortex. Science 277, 968–971 (1997).

10. Johansen, J. P., Fields, H. L. & Manning, B. H. The affective component of pain in rodents: direct evidence for a contribution of the anterior cingulate cortex. Proc Natl Acad Sci U S A 98, 8077–8082 (2001).

11. Johansen, J. P. & Fields, H. L. Glutamatergic activation of anterior cingulate cortex produces an aversive teaching signal. Nat Neurosci 7, 398–403 (2004).

12. Zhang, Q. et al. Chronic pain induces generalized enhancement of aversion. eLife 6, e25302 (2017).

13. Singh, A. et al. Mapping Cortical Integration of Sensory and Affective Pain Pathways. Curr Biol 30, 1703–1715.e5 (2020).

14. Li, X.-Y. et al. Alleviating neuropathic pain hypersensitivity by inhibiting PKMzeta in the anterior cingulate cortex. Science 330, 1400–1404 (2010).

15. Zhou, H. et al. Ketamine reduces aversion in rodent pain models by suppressing hyperactivity of the anterior cingulate cortex. Nat Commun 9, 3751 (2018).

16. Golden, C. T. & Chadderton, P. Psilocybin reduces low frequency oscillatory power and neuronal phase-locking in the anterior cingulate cortex of awake rodents. Sci Rep 12, 12702 (2022).

17. Kim, K. et al. Structure of a Hallucinogen-Activated Gq-Coupled 5-HT2A Serotonin Receptor. Cell 182, 1574–1588.e19 (2020).

18. Gregory, N. S. et al. No evidence of immediate or persistent analgesic effect from a single dose of psilocybin in three mouse models of pain. Nat Commun 17, 1916 (2026).

19. Singh, A. et al. Mapping Cortical Integration of Sensory and Affective Pain Pathways. Current Biology 30, 1703–1715.e5 (2020).

20. Dale, J. et al. Scaling Up Cortical Control Inhibits Pain. Cell Rep 23, 1301–1313 (2018).

21. Zhou, H. et al. Ketamine reduces aversion in rodent pain models by suppressing hyperactivity of the anterior cingulate cortex. Nat Commun 9, 3751 (2018).

22. King, T. et al. Unmasking the tonic-aversive state in neuropathic pain. Nat Neurosci 12, 1364–1366 (2009).

23. Johansen, J. P., Fields, H. L. & Manning, B. H. The affective component of pain in rodents: direct evidence for a contribution of the anterior cingulate cortex. Proc Natl Acad Sci U S A 98, 8077–8082 (2001).

24. Corne, S. J. & Pickering, R. W. A possible correlation between drug-induced hallucinations in man and a behavioural response in mice. Psychopharmacologia 11, 65–78 (1967).

25. Bedard, P. & Pycock, C. J. ‘Wet-dog’ shake behaviour in the rat: a possible quantitative model of central 5-hydroxytryptamine activity. Neuropharmacology 16, 663–670 (1977).

26. Halberstadt, A. L. & Geyer, M. A. Multiple receptors contribute to the behavioral effects of indoleamine hallucinogens. Neuropharmacology 61, 364–381 (2011).

27. Bouloufa, A. et al. LSD’s rapid antidepressant effects are modulated by 5-HT2B receptors. Biomedicine & Pharmacotherapy 190, 118348 (2025).

28. Vetulani, J., Bednarczyk, B., Reichenberg, K. & Rokosz, A. Head twitches induced by LSD and quipazine: Similarities and differences. Neuropharmacology 19, 155–158 (1980).

29. Xiao, Z. et al. Cortical Pain Processing in the Rat Anterior Cingulate Cortex and Primary Somatosensory Cortex. Front. Cell. Neurosci. 13, (2019).

30. Rainville, P., Duncan, G. H., Price, D. D., Carrier, B. & Bushnell, M. C. Pain affect encoded in human anterior cingulate but not somatosensory cortex. Science 277, 968–971 (1997).

31. Araneda, R. & Andrade, R. 5-Hydroxytryptamine2 and 5-hydroxytryptamine 1A receptors mediate opposing responses on membrane excitability in rat association cortex. Neuroscience 40, 399–412 (1991).

32. Schmitz, G. P. et al. Psychedelic compounds directly excite 5-HT2A layer V medial prefrontal cortex neurons through 5-HT2A Gq activation. Transl Psychiatry 15, 381 (2025).

33. Aghajanian, G. K. & Marek, G. J. Serotonin, via 5-HT2A receptors, increases EPSCs in layer V pyramidal cells of prefrontal cortex by an asynchronous mode of glutamate release. Brain Res 825, 161–171 (1999).

34. Shao, L.-X. et al. Psilocybin induces rapid and persistent growth of dendritic spines in frontal cortex in vivo. Neuron 0, (2021).

35. Brys, I. et al. 5-HT2AR and NMDAR psychedelics induce similar hyper-synchronous states in the rat cognitive-limbic cortex-basal ganglia system. Commun Biol 6, 737 (2023).

36. Wood, J., Kim, Y. & Moghaddam, B. Disruption of Prefrontal Cortex Large Scale Neuronal Activity by Different Classes of Psychotomimetic Drugs. J. Neurosci. 32, 3022–3031 (2012).

37. Dearnley, B., Dervinis, M., Shaw, M. & Okun, M. Stretching and squeezing of neuronal log firing rate distribution by psychedelic and intrinsic brain state transitions. 2021.08.22.457198 Preprint at 10.1101/2021.08.22.457198 (2021).

38. Dale, J. et al. Scaling Up Cortical Control Inhibits Pain. Cell Rep 23, 1301–1313 (2018).

