## Supplemental Figures for "LSD persistently disrupts affective pain processing"

### Supplementary figures and legends

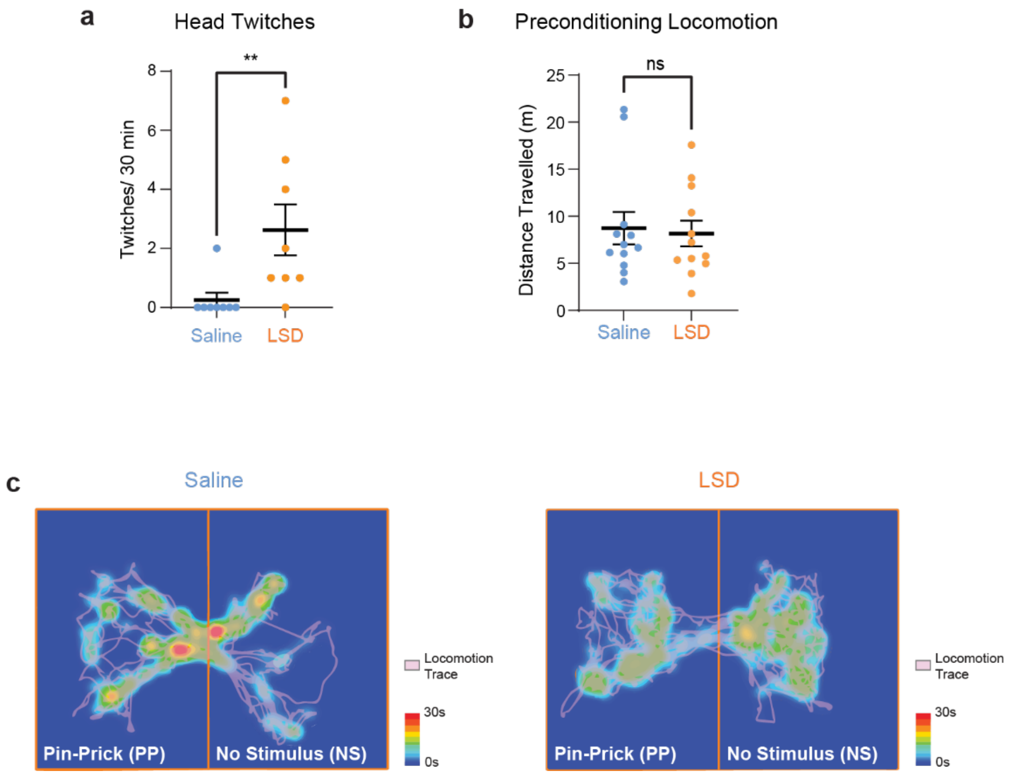


#### Supplementary Figure 1

a. Rats given IP LSD engage in significantly more head twitches than those given IP saline (saline 0.25 ± 0.25 twitches/ 30 min, n=8, LSD 2.62 ± 0.86 twitches/ 30 min, unpaired Student's t-test: $p = 0.006$, n = 8).

b. There is no difference between locomotion between rats given IP saline vs. IP LSD (Saline 8.73 ± 1.72, n=12; LSD mean = 8.16 ± 1.36, n=12; unpaired Student's t-test: $p = 0.80$).

c. Left: Locomotion heatmap of a rat given IP saline during the pre-conditioning phase. Right: Locomotion heatmap of a rat given IP LSD during the pre-conditioning phase.

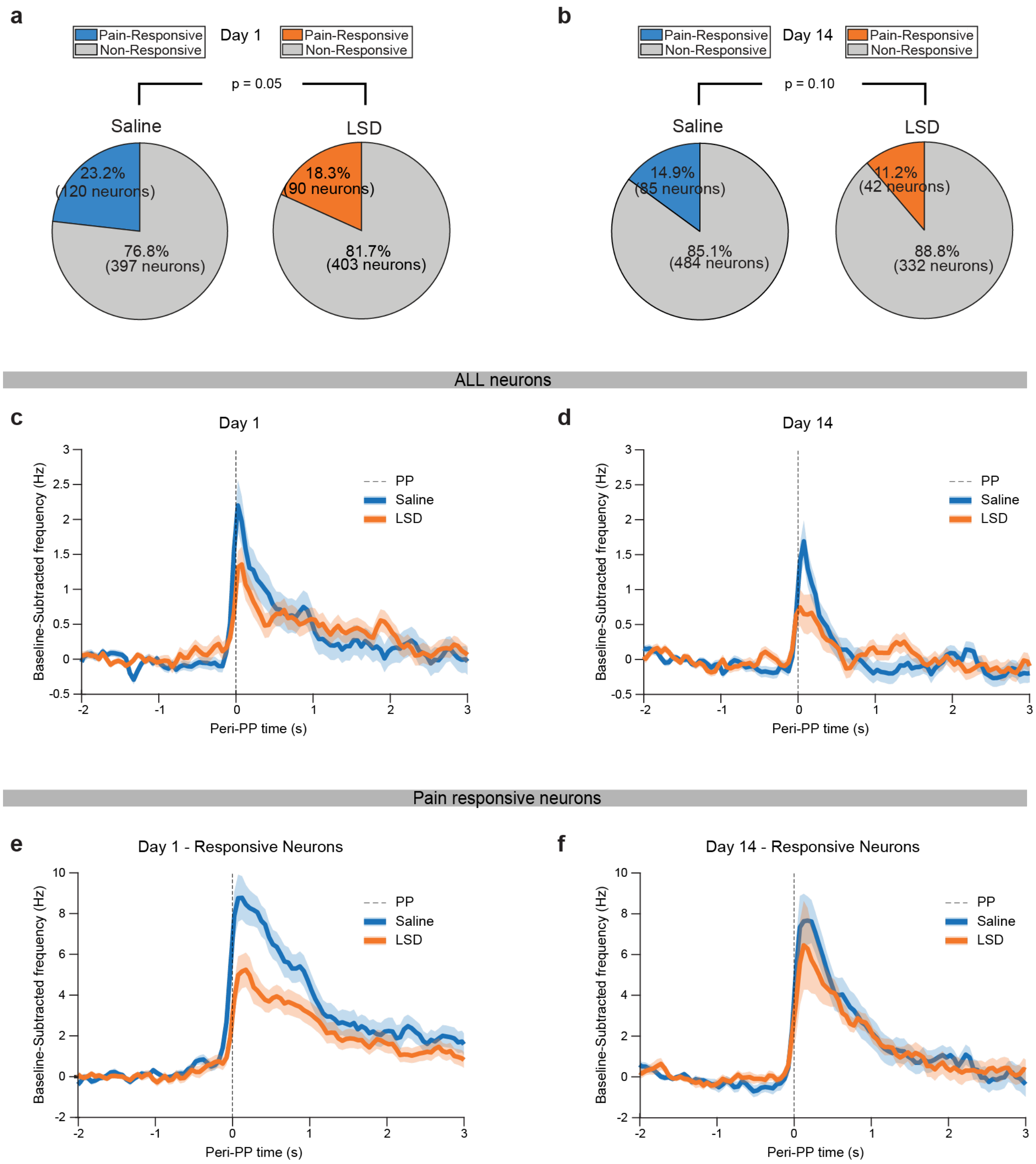


### Supplementary Figure 2

a. Percentage of pain responsive neurons in rats on day 1 of injection (Chi square: $p = 0.05$).

b. Percentage of pain responsive neurons in rats on day 14 of injection (Chi square: $p = 0.10$).

c. Traces of neuronal firing rate following pin prick administration for neurons 15 minutes after exposure to saline (N = 4 rats, 515 neurons) or LSD (N = 4 rats, 486 neurons).

d. Traces of neuronal firing rate following pin prick administration for neurons 14 days after exposure to saline (N = 4 rats, 569 neurons) or LSD (N = 4 rats, 372 neurons).

e. Traces of neuronal firing rate following pin prick administration for pain-responsive neurons 15 minutes after exposure to saline (N = 4 rats, 120 neurons) or LSD (N = 4 rats, 90 neurons).

f. Traces of neuronal firing rate following pin prick administration for pain-responsive neurons 14 days after exposure to saline (N = 4 rats, 85 neurons) or LSD (N = 4 rats, 42 neurons).

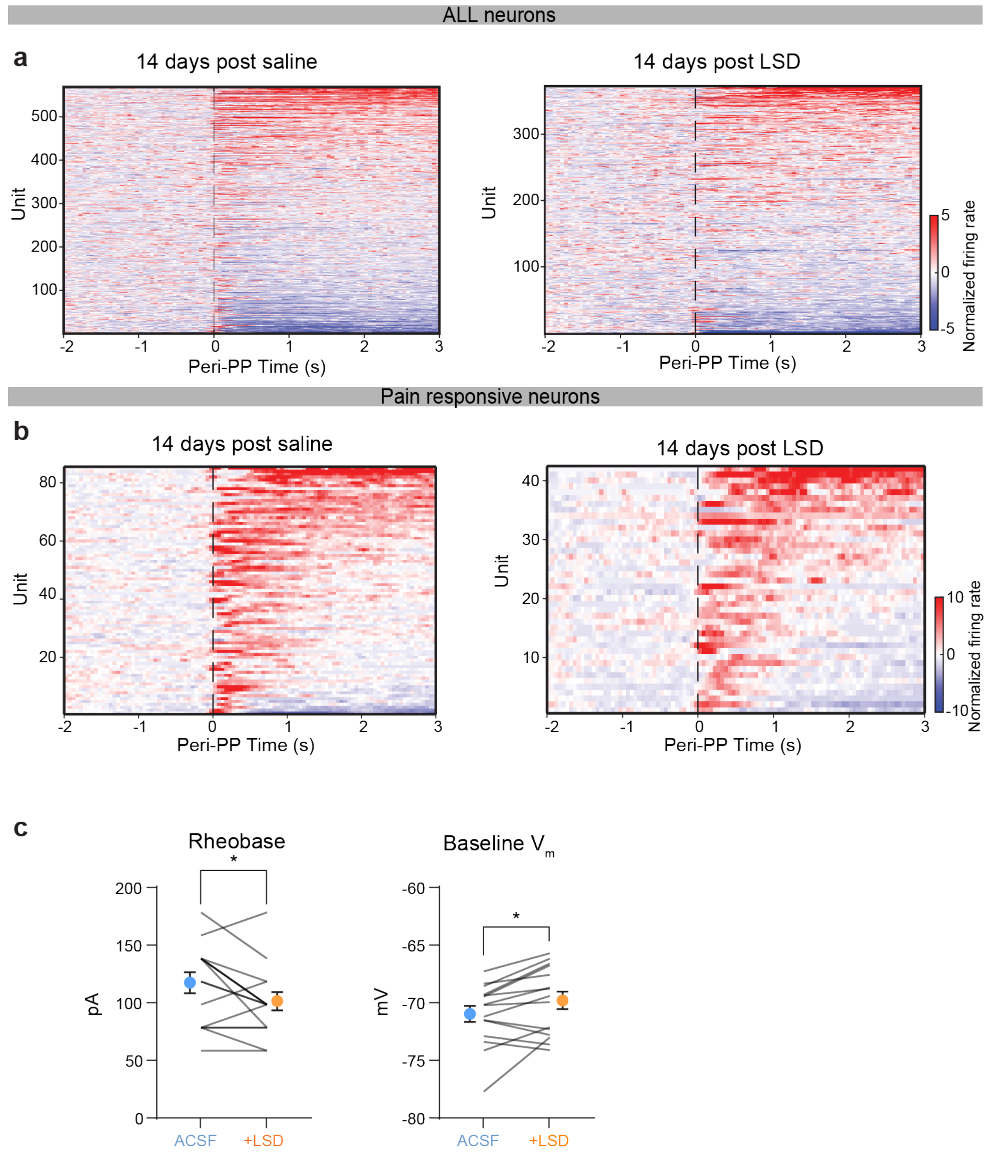


### Supplementary Figure 3

a. Left: heatmap of all neurons 14 days after exposure to saline. Pin prick is applied at time 0. Firing rate normalized relative to pre pin prick activity (N = 4 rats, 569 neurons). Right: heatmap of all neurons 14 days after exposure to LSD. Pin prick is applied at time 0. Firing rate normalized relative to pre pin prick activity (N = 4 rats, 372 neurons).

b. Left: heatmap of pain-responsive neurons 14 days after exposure to saline. Pin prick is applied at time 0. Firing rate normalized relative to pre pin prick activity (N = 4 rats, 85 neurons). Right: heatmap of all neurons 14 days after exposure to saline. Pin prick is applied at time 0. Firing rate normalized relative to pre pin prick activity (N = 4 rats, 42 neurons).

c. Left: Neurons exposed to LSD had significantly lower rheobase ($n = 15$ neurons; Wilcoxon paired test: $p = 0.03$). Right: Neurons exposed to LSD had significantly higher baseline membrane potential ($V_m$; $n = 15$ neurons; paired student's t-test: $p = 0.02$).
